# Visual Perception in the Periphery: The Role of Covert Attention Vectors in the Extraction of Semantic Information

**DOI:** 10.1101/2020.08.02.231803

**Authors:** Ikaasa Suri, Patrick McGranor Wilson, Saba Doustmohammadi, Anna De Schutter, Thida Sandy Chunwatanapong, Juanyi Tan, Sara Divija Varadharajulu, Nicholas Hunter O’Connell, Archibald Lai, Sakshi Dureja, River Jonathan Phoenix Govin, Katsushi Arisaka, Elizabeth Anne Falcone Mills

**Author notes:** Both Co-first authors contributed equally to this paper (Ikaasa Suri, Patrick McGranor Wilson). Corresponding Author: Elizabeth Anne Falcone Mills, Ph.D.

## Abstract

Under covert attention, our visual perception deteriorates dramatically as eccentricity increases. This reduction of peripheral visual acuity (PVA) is partially due to the coarse sampling of the retinal ganglion cells towards the periphery, but this property cannot be solely responsible. Other factors, such as character crowding, have been studied, yet the origin of the poor PVA is not entirely understood. This gap motivated us to investigate the PVA by varying the crowding conditions systematically. Under completely crowded conditions (i.e. resembling a full page of text), PVA was observed to be eight times worse than the PVA under uncrowded conditions. By partially crowding the periphery, we obtained PVA values between the fully crowded and uncrowded conditions. On the other hand, crowding the fovea center while leaving the periphery uncrowded improved PVA relative to the uncrowded case. These results support a model for a top-down “covert attention vector” that assists the resulting PVA in a manner analogous to saccadic eye movement for overt attention. We speculate that the attention vector instructs the dorsal pathway to transform the peripheral character to the foveal center. Then, the scale-invariant log-polar retinotopy of the ventral pathway can scale the centered visual input to match the prior memory of the specific character shape.

## Introduction

The semantic extraction of visual information is remarkable: whether shifted, rotated, or scaled, we are still able to perceive the identity of an object (Baars & Gage, 2010; Hebb, 1949). While the mechanism behind peripheral visual acuity (PVA) has not been fully elucidated, prior work has suggested several factors that impair semantic extraction in the periphery. Most important among these factors are the increase in the peripheral retinal ganglion cell receptive field size and the decrease in cortical magnification factor (i.e. the number of neurons in the visual cortex that are responsible for processing a given visual stimulus) (Harvey & Dumoulin, 2011; Horton & Hoyt, 1991; McCann, Hayhoe, & Geisler, 2011; Wienbar & Schwartz, 2018).

However, these findings are not sufficient to explain the entire mechanism behind semantic extraction in the periphery: while PVA is generally considered imprecise and relatively inaccurate compared to visual acuity at the fovea, there exist certain conditions under which PVA can be better (Anstis, 1974; Lettvin, 1976; Strasburger, Rentschler, & Jüttner, 2011). S.M. Anstis (1974) initially observed that threshold letter height (i.e. minimum distinguishable letter size) increases 0.05° for every one-degree increase in retinal eccentricity (i.e. the angle between the fovea and the visual stimulus).

Research on character crowding has attempted to explain the factors that impact PVA under various conditions. In particular, crowding has been defined as the distance between the center of the flanks (i.e. distractors) and the center of the visual stimulus (i.e. the object being identified) (Levi & Carney, 2009; Pelli, 2008). In both the periphery and, to a lesser extent, at the fovea, crowding has been shown to worsen visual acuity (Bouma, 1970; Lalor, Formankiewicz, & Waugh, 2016; Rosen, Chakravarthi, & Pelli, 2014; Yehezkel et al., 2015). However, the majority of this past work has studied crowding only in the sense that it either impairs visual acuity if it is present or does not impair visual acuity if the critical spacing threshold is not met; from this work, we have been unable to determine the mechanism by which crowding impedes visual acuity (Levi, 2011).

While the retinotopic mapping of visual space in the ventral pathway has not been discussed within the context of PVA, its log-polar coordinate system may provide some insight into how we are able to extract semantic information in the periphery. Previous research concluded that visuospatial information is mapped onto the primary visual cortex (V1) in a log-polar manner (Abdollahi et al., 2014; Benson et al., 2014; Horton & Hoyt, 1991; Wandell, Dumoulin, & Brewer, 2007). However, studies within the vision community have not discussed the way in which this log-polar mapping system can be utilized for scaling invariant perception. While literature on computer vision has acknowledged the role of scale invariance in vision, it has not yet been applied to human vision (Araujo & Dias, 1997; Qi, Bao, & Nakata, 2006; Traver & Bernardino, 2010). After the input is transformed into a log-polar image on the V1, it maintains the same spatial configuration independent of scaling in visual space; with this log-polar coordinate system, scaling is represented by translation (Appendix Figure 1).

It is with this concept of scaling-invariant perception that we hypothesize the existence of an attentional, parallel-process model for visual perception that allows for top-down communication (Fries, 2015; Friston et al., 2018; Gilbert & Li, 2013; Klimesch, 2012). Specifically, because it is improbable that we have an infinite number of semantic memories for a single object, we propose that our brain utilizes a model by which we retain one semantic memory for each object. In order for us to recognize an object presented to us in visual space, our input must be matched against this prior pattern through coincidence detection (DiCarlo, Zoccolan, & Rust, 2012; Kaiser et al. 2019). Thus, no matter the location of the object––in our periphery or at the fovea––or its size, we are able to generate the same semantic sensation when it is in our visual field. Each time a single object is recognized––that is, each time our semantic memory and the visual input overlap––we hypothesize that the pathway is reinforced through Hebbian plasticity (Hebb, 1949). Given that visual acuity is the greatest at the fovea, this prior pattern must be the neuronal pattern that is also activated when an object is perceived at the fovea (Appendix Figure 2). However, due to this same log-polar mapping system, visual input from the periphery is significantly distorted relative to the foveal image that our brain has been trained to recognize. Thus, current work does not explain the mechanism through which we are able to process peripheral semantic information.

Importantly, prior studies were only interested in determining the critical spacing between characters that is necessary for the crowding effect to impact visual acuity. Building off of this work, we examine crowding not as a function of critical spacing between characters but instead as a function of its location––at the fovea or in the periphery––and its extent. Under these varied conditions, we aim to better understand the mechanism of semantic extraction from peripheral visual input. We created a series of protocols that examine threshold letter height at set retinal eccentricities. In particular, we investigate (1) PVA as a function of eccentricity, (2) the extent to which a greater amount of crowding impedes PVA, and (3) whether crowding at the fovea impacts PVA. These data were then analyzed to provide further insight into how peripheral semantic information is extracted.

## Methods

### Ethics Statement

This study was approved by the University of California, Los Angeles Institutional Review Board (IRB#19-000751). All recruited subjects were volunteers who provided two forms of consent: (1) written consent prior to scheduling a study session and (2) oral consent prior to performing the protocols.

### Procedure Summary

Subjects performed all protocols in a dark room that was illuminated solely by the monitor displaying the paradigm. These lumination settings were sufficient in order for the subjects to utilize photopic vision. The subjects rested their chin on a chinrest placed 50 cm away from the monitor that was adjusted to the subjects’ height. The chinrest stabilized the head movement of the subject throughout the study session (Figure 1).

**Figure 1.**
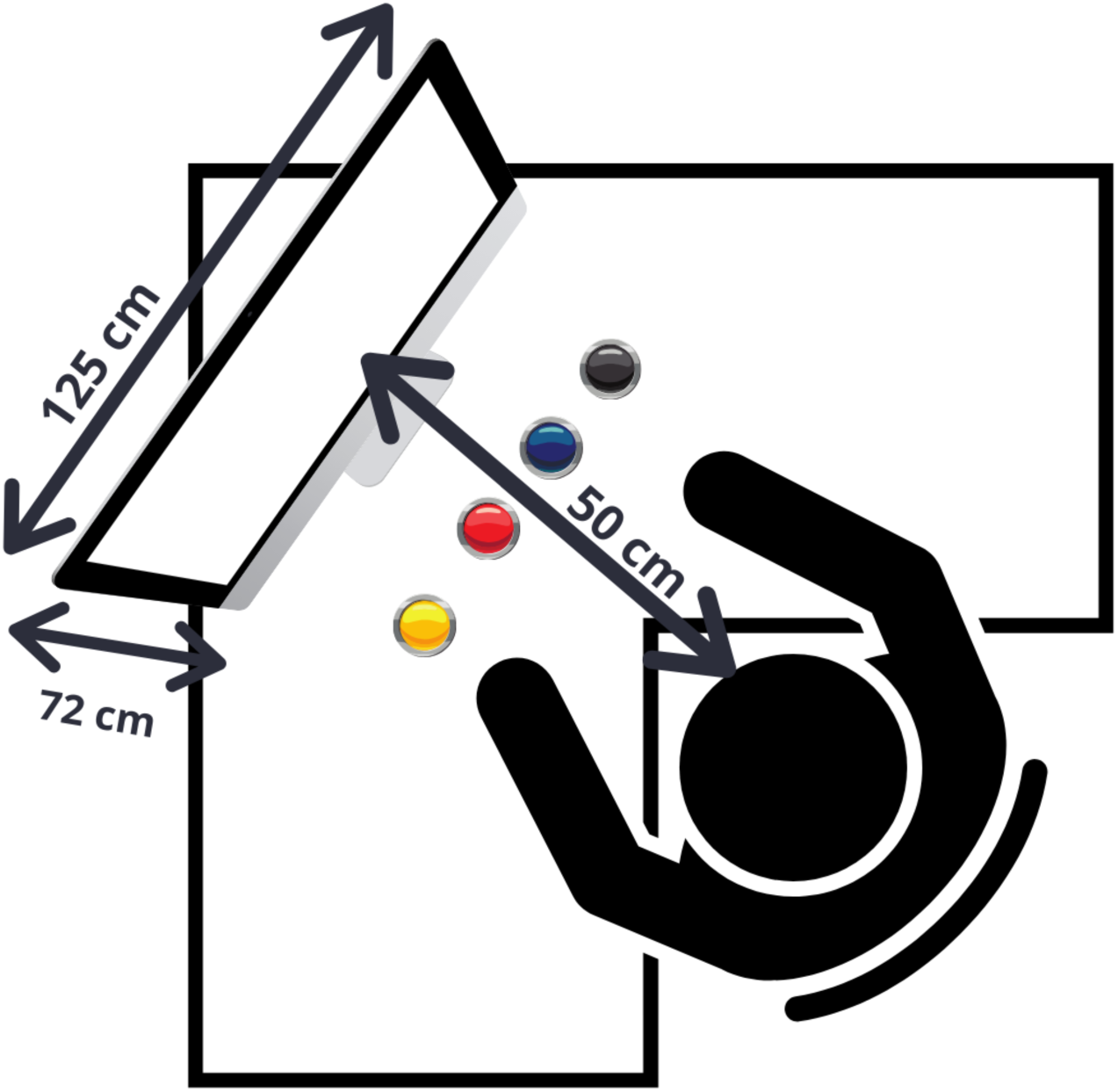
Bird’s eye view of study session set-up for subjects. A 55 inch TV with a viewable display size of 138.7 cm (54.6 in) and a panel resolution of 3840 x 2160 pixels was used to display paradigms (4k / 60 Hz, TCL, Model 55S424). Subject input was recorded through a physical push button setup built on a breadboard and run by an Arduino UNO Rev 3 (ELEGOO, Shenzhen, China), which interfaced with MATLAB and Python via serial output. Subjects had the option of pressing one of four buttons during each trial: (1) a yellow button for “B”, (2) a red button for “E”, (3) a blue button for “P”, or (4) a black button for “I don’t know”.

A 55 inch TV with a viewable display size of 138.7 cm (54.6 in) was used to display stimuli (4k / 60 Hz, TCL, Model 55S424). Stimuli for the fully crowded protocol were generated in MATLAB and displayed using the Psychophysics Toolbox (Brainard, 1997; Pelli, 1997). Stimuli for all other protocols were generated in Python 3 and displayed using the PsychoPy library (Pierce et al., 2019).

Each protocol, described in more detail under *Behavioral Tasks*, consisted of a series of white letters displayed upon a grey background. The letters appeared at some position in the subject’s periphery, and the subject was asked to identify the stimulus letter during each trial. A small green dot appeared in the center of the screen at the beginning of each protocol, and subjects were instructed to fix their gaze upon it for the duration of each protocol. All letters displayed throughout the protocols were randomly selected from a pool of three capital letters: B, E, and P. These letters were selected because prior literature suggests that, due to their visual similarities, precise semantic recognition is required to differentiate between them (Mathew, Shah, & Simon, 2011).

Subject input was recorded through a physical push button setup built on a breadboard and run by an Arduino UNO Rev 3 (ELEGOO, Shenzhen, China), which interfaced with MATLAB and Python via serial output. Subjects had the option of pressing one of four buttons during each trial: (1) a yellow button for “B”, (2) a red button for “E”, (3) a blue button for “P”, or (4) a black button for “I don’t know”. Subjects pressed the button corresponding to the letter perceived in their periphery. They pressed the black button only when they were entirely unable to discern which character was displayed in the periphery or if they shifted their gaze from the centralized dot. The purpose of this black button was to reduce the potential for lucky guessing and account for a breach in protocol.

For the Fully Crowded and Three Lines protocols, subjects continually identified stimulus characters along the horizontal axis. The trial for these two protocols continued until a stimulus character was identified incorrectly. For all other protocols, the subject continued to record their response via the buttons until a staircase algorithm determined the threshold letter height at a particular radius away from the center of the screen (i.e. at a set retinal eccentricity). In these protocols, a correct identification of the character resulted in a new randomly selected character being displayed with a slight reduction in the letter height, whereas an incorrect identification resulted in a slight increase. A “reversal” occurred when an incorrect response followed a correct response, or a correct response followed an incorrect response. The trial continued until three consecutive reversals were achieved, after which the staircase algorithm would restart at a different retinal eccentricity.

At the end of each trial for all protocols, the direction, threshold letter height, and retinal eccentricity in degrees were recorded. Retinal eccentricity was defined as the angle subtended by the distance from the subject’s retina to the screen and the distance from the stimulus character to the screen’s center. Similarly, letter height was defined as the angle subtended by the distance from the subject’s retina to the screen and the distance from the top to the bottom of the stimulus character. The stimuli in the Fully Crowded protocol were displayed using Helvetica font; stimuli in all other protocols were displayed in Arial font. The source code for all of the behavioral projects is publicly available on GitHub (Wilson, 2020).

### Behavioral Tasks

#### Protocol 1: Fully Crowded (Fig. 2a)

The stimulus consisted of a block of letters of a given height which covered the entirety of the display. Subjects were instructed to read in a straight line away from the center dot on the horizontal axis. The direction in which the subject was required to read for each trial was indicated by a small arrow in the center of the green dot. Upon incorrectly identifying the character, a new letter height was randomly selected and a new randomized block of text was generated. The subject repeated this for a letter height of 0.23° and all letter heights in the range of 0.5° to 4° in increments of 0.5°, and 5° to 10° in increments of 1°.

**Figure 2.**
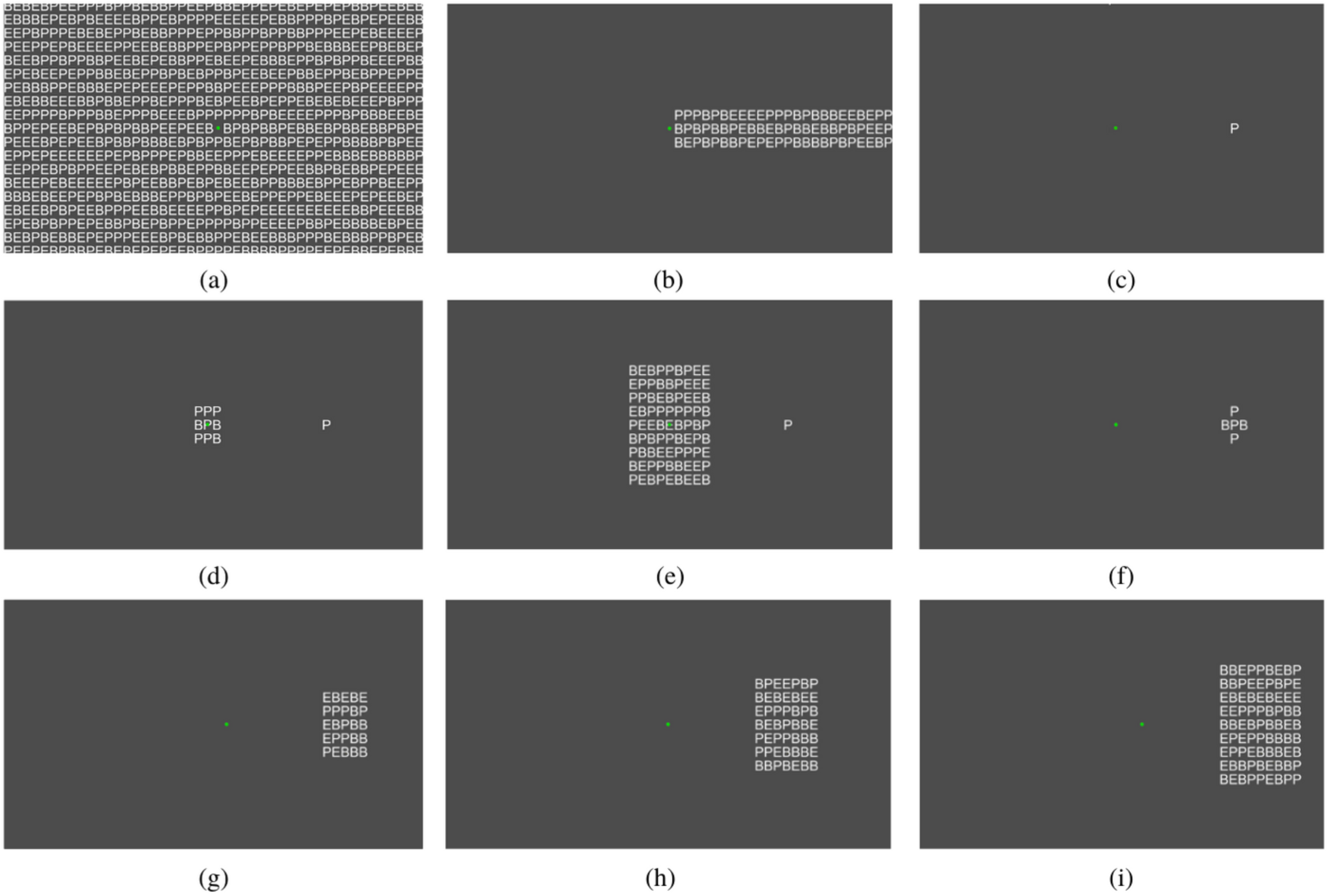
Behavioral paradigm displays. A green dot was centered on the screen for each paradigm. Subjects were asked to fix their gaze upon the green dot for the duration of the paradigm. (a) Fully Crowded, (b) Three Lines, (c) Isolated Character, (d) Crowded Center 3×3, (e) Crowded Center 9×9, (f) Crowded Periphery Center, Crowded Periphery Inner, and Crowded Periphery Outer, (g) Crowded Periphery 5×5, (h) Crowded Periphery 7×7, (i) Crowded Periphery 9×9.

#### Protocol 2: Three Lines (Fig. 2b)

The stimulus consisted of three horizontal lines of letters of a given height. Subjects were instructed to read the characters on the middle horizontal line, moving from the center green dot to the periphery. The three lines only appeared in one direction (i.e. either to the left or to the right of the green dot). Upon incorrectly identifying a character, a new letter height was randomly selected, and a new set of letters for the three lines was generated. The subject repeated this task for all letter heights in the range of 0.5° to 4°, in increments of 0.5°.

#### Protocol 3: Isolated Character (Fig. 2c)

The target stimulus consisted of a single letter. The stimulus character was displayed at a randomly selected retinal eccentricity in the range of 0° to 40° on the horizontal axis. The trial continued until the aforementioned staircase algorithm determined the threshold letter height. At the start of the next trial, a new retinal eccentricity was randomly selected. The task was repeated for all retinal eccentricities in the range of 0° to 40°, in increments of 5°.

#### Protocol 4: Crowded Center (Fig. 2d and 2e)

The target stimulus consisted of a single letter. A square block of white letters was also displayed at the center of the screen, all of which were displayed at the same letter height as the stimulus character in the periphery. The stimulus character was displayed at a randomly selected retinal eccentricity on the horizontal axis. Each time the subject recorded a response, a new block of text in the center of the screen and a new stimulus character in the periphery were randomly generated. The trial continued until the aforementioned staircase algorithm determined the threshold letter height. At the start of the next trial, a new retinal eccentricity was randomly selected. The task was repeated for all retinal eccentricities in the range of 10° to 40°, in increments of 5°.

There were two variations of this protocol. In Crowded Center 3×3 (Fig. 2d), a 3 by 3 block of text was displayed in the center of the screen. In Crowded Center 9×9 (Fig. 2e), a 9 by 9 block of text was displayed in the center of the screen.

#### Protocol 4: Crowded Periphery – “Plus” (Fig. 2f)

The stimulus consisted of a “plus” pattern of letters. The stimulus was displayed at a randomly selected retinal eccentricity in the range of 5° to 40° on the horizontal axis. Each time the subject recorded a response, a new peripheral stimulus was randomly generated. The task was repeated for all retinal eccentricities in the range of 5° to 40°, in increments of 5°.

There were three variations of this protocol. While the display remained the same, the task was different. For the Crowded Periphery Inner protocol, the subject was asked to identify the character on the inside of the “plus” pattern (i.e. closest to the green dot). For the Crowded Periphery Center protocol, the subject was asked to identify the character in the center of the “plus” pattern. For the Crowded Periphery Outer protocol, the subject was asked to identify the character on the outside of the “plus” pattern (i.e. furthest from the green dot).

#### Protocol 5: Crowded Periphery – “Block” (Fig. 2g-2i)

The stimulus consisted of a block of letters. The stimulus was displayed at a randomly selected retinal eccentricity in the range of 5° to 40° on the horizontal axis. Each time the subject recorded a response, a new peripheral stimulus was randomly generated. The task was repeated for all retinal eccentricities in the range of 5° to 40°, in increments of 5°.

There were three variations of this protocol. In Crowded Periphery 5×5 (Fig. 2g), the stimulus character to be identified was the character at the center of a 5×5 block of text; similarly, in Crowded Periphery 7×7 (Fig. 2h) and Crowded Periphery 9×9 (Fig. 2i), the stimulus character was at the center of a 7×7 block of text and a 9×9 block of text, respectively.

### Subjects

Two pools of subjects participated: (1) subjects recruited from the general UCLA student population, per IRB-approved procedures and (2) internal researchers who collected pilot data by performing the protocols in mock study sessions. In total, there were 17 data sets for the Fully Crowded protocol, 3 data sets for the Three Lines protocol, 21 data sets for the Isolated Character protocol, 6 data sets for the Crowded Center 3×3 protocol, 5 data sets for the Crowded Center 9×9 protocol, 9 data sets for the Crowded Periphery Center protocol, 9 data sets for the Crowded Periphery Outer protocol, 5 data sets for the Crowded Periphery 5×5 protocol, 5 data sets for the Crowded Periphery 7×7 protocol, and 4 data sets for the Crowded Periphery 9×9 protocol. Table 2 provides a more detailed subject breakdown.

**Table 1.** Summary comparing stimulus and analysis details across behavioral protocols. X indicates the presence of a given parameter. In Crowded Periphery (CP) visual acuity paradigms, extra (distracting) letters flanked the peripheral target character. In Crowded Center (CC) visual acuity paradigms, extra (distracting) letters flanked the center green dot. The use of a staircase algorithm provided increased accuracy in finding the threshold letter height for each eccentricity location tested. Stimulus characters were displayed in the Arial font in all protocols except for Fully Crowded, which used the Helvetica font.

| *Visual Acuity Paradigm* | *Protocol Parameters* |  |  |  |
| --- | --- | --- | --- | --- |
|  | **Peripheral Crowding** | **Center Crowding** | **Use of Staircase Algorithm** | **Font** |
| Fully Crowded (FC) | **X** | **X** |  | *Helvetica* |
| Three Lines (3L) | **X** | **X** |  | *Arial* |
| Isolated Character (IC) |  |  | **X** | *Arial* |
| **Crowded Center (CC)** |  |  |  |  |
| Crowded Center 3x3 (CC3) |  | **X** | **X** | *Arial* |
| Crowded Center 9x9 (CC9) |  | **X** | **X** | *Arial* |
| **Crowded Periphery (CP)** |  |  |  |  |
| Crowded Periphery Inner (CPI) | **X** |  | **X** | *Arial* |
| Crowded Periphery Center (CPC) | **X** |  | **X** | *Arial* |
| Crowded Periphery Outer (CPO) | **X** |  | **X** | *Arial* |
| Crowded Periphery 5x5 (CP5) | **X** |  | **X** | *Arial* |
| Crowded Periphery 7x7 (CP7) | **X** |  | **X** | *Arial* |
| Crowded Periphery 9x9 (CP9) | **X** |  | **X** | *Arial* |

**Table 2.**
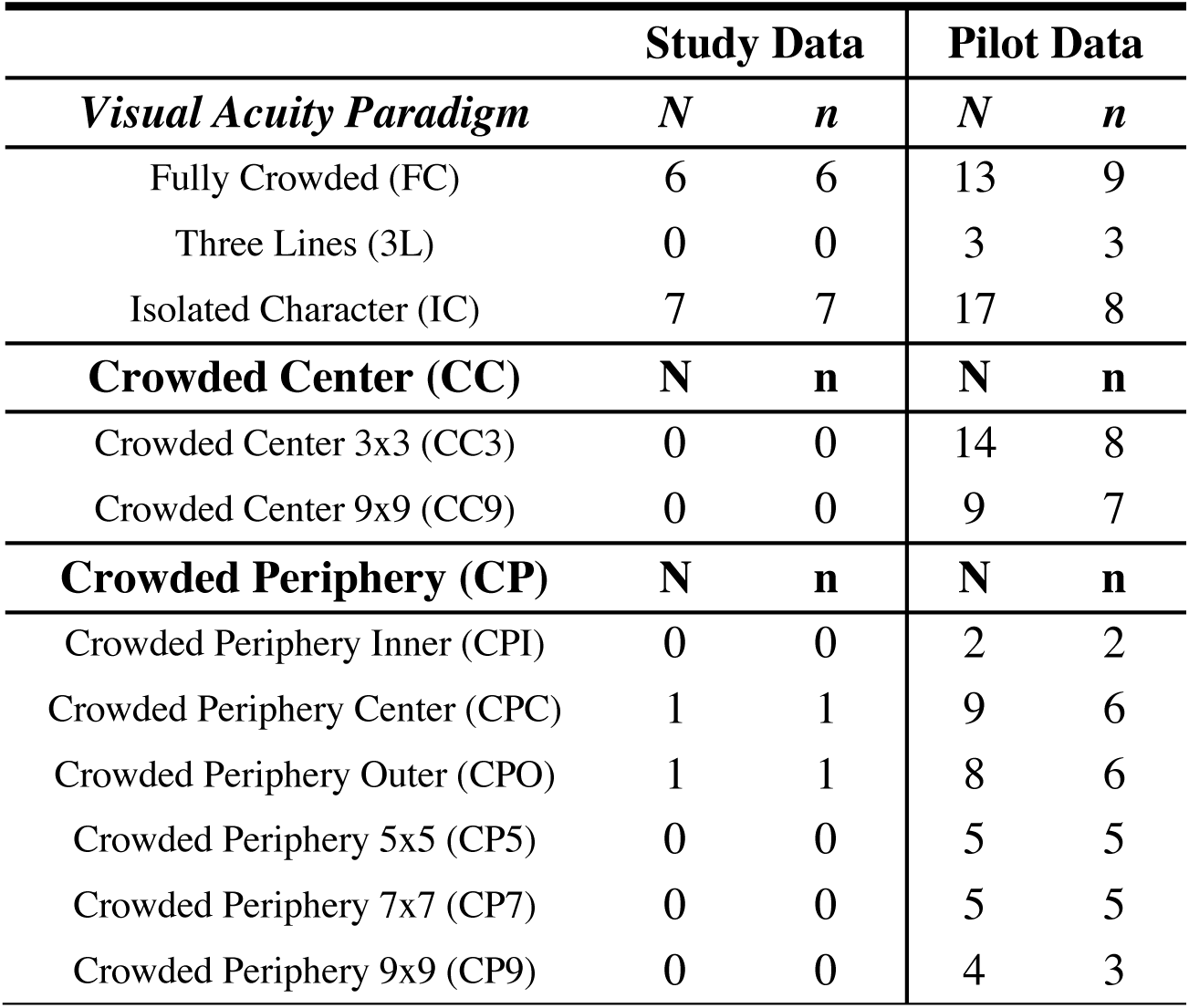
Behavioral protocol breakdown according to subject pool. *N* refers to the number of individual data sets for the specified paradigm; *n* refers to the number of individual subjects who performed the specified paradigm. Subjects were split into two pools: (1) study data (i.e. subjects recruited from the general UCLA student population, per IRB-approved procedures) and (2) pilot data (i.e. internal researchers who collected pilot data by performing the protocols in mock study sessions).

|  | **Study Data** |  | **Pilot Data** |  |
| --- | --- | --- | --- | --- |
| ***Visual Acuity Paradigm*** | **$N$** | **$n$** | **$N$** | **$n$** |
| Fully Crowded (FC) | 6 | 6 | 13 | 9 |
| Three Lines (3L) | 0 | 0 | 3 | 3 |
| Isolated Character (IC) | 7 | 7 | 17 | 8 |
| **Crowded Center (CC)** | **$N$** | **$n$** | **$N$** | **$n$** |
| Crowded Center 3x3 (CC3) | 0 | 0 | 14 | 8 |
| Crowded Center 9x9 (CC9) | 0 | 0 | 9 | 7 |
| **Crowded Periphery (CP)** | **$N$** | **$n$** | **$N$** | **$n$** |
| Crowded Periphery Inner (CPI) | 0 | 0 | 2 | 2 |
| Crowded Periphery Center (CPC) | 1 | 1 | 9 | 6 |
| Crowded Periphery Outer (CPO) | 1 | 1 | 8 | 6 |
| Crowded Periphery 5x5 (CP5) | 0 | 0 | 5 | 5 |
| Crowded Periphery 7x7 (CP7) | 0 | 0 | 5 | 5 |
| Crowded Periphery 9x9 (CP9) | 0 | 0 | 4 | 3 |

### Statistical Analysis

Threshold letter height was plotted against retinal eccentricity. A line of best fit was calculated for each protocol via *χ*^2^ minimization and validated by examination of the *χ*_*ν*_^2^ statistic. Each subject’s slope parameter from every protocol was plotted against their Isolated Character slope parameter to assess correlation between protocol performances. Data across all subjects were aggregated and threshold letter height observations were normalized by dividing by their respective retinal eccentricity values, *Threshold Letter Height/Retinal Eccentricity*. Significance tests were performed upon the resulting distribution to assess the effects of the various protocols upon visual acuity. Significance tests were also performed on the distributions of measurements taken in the left vs. right visual fields. A detailed explanation of the statistical analysis conducted can be found in the appendix.

## Results

We observed a spectrum of visual acuity that was dependent on the crowding both in the periphery and at the center gaze location. This method of analysis allowed for key insights into the data. More specifically, we were able to discern two broadly categorized groups within our dataset: (1) those that support previous literature and (2) those that are novel with regards to PVA. In particular, we focus on the relationship between the Fully Crowded and the Crowded Periphery 9×9 protocols, the Isolated Character and Crowded Center protocols, and the Crowded Periphery Inner and Crowded Periphery Outer protocols.

### Letter Height vs. Retinal Eccentricity

Visual acuity data collected from the right visual field (i.e. periphery to the right of the center green dot) and from the left visual field (i.e. periphery to the left of the center green dot) were lumped and plotted for each protocol (Figure 3). Visual acuity for stimuli displayed in the right visual field was better than for stimuli displayed in the left visual field in the Isolated Character paradigm (p = 0.009). There was no significance observed between the left and right visual fields for any other protocols.

**Figure 3.**
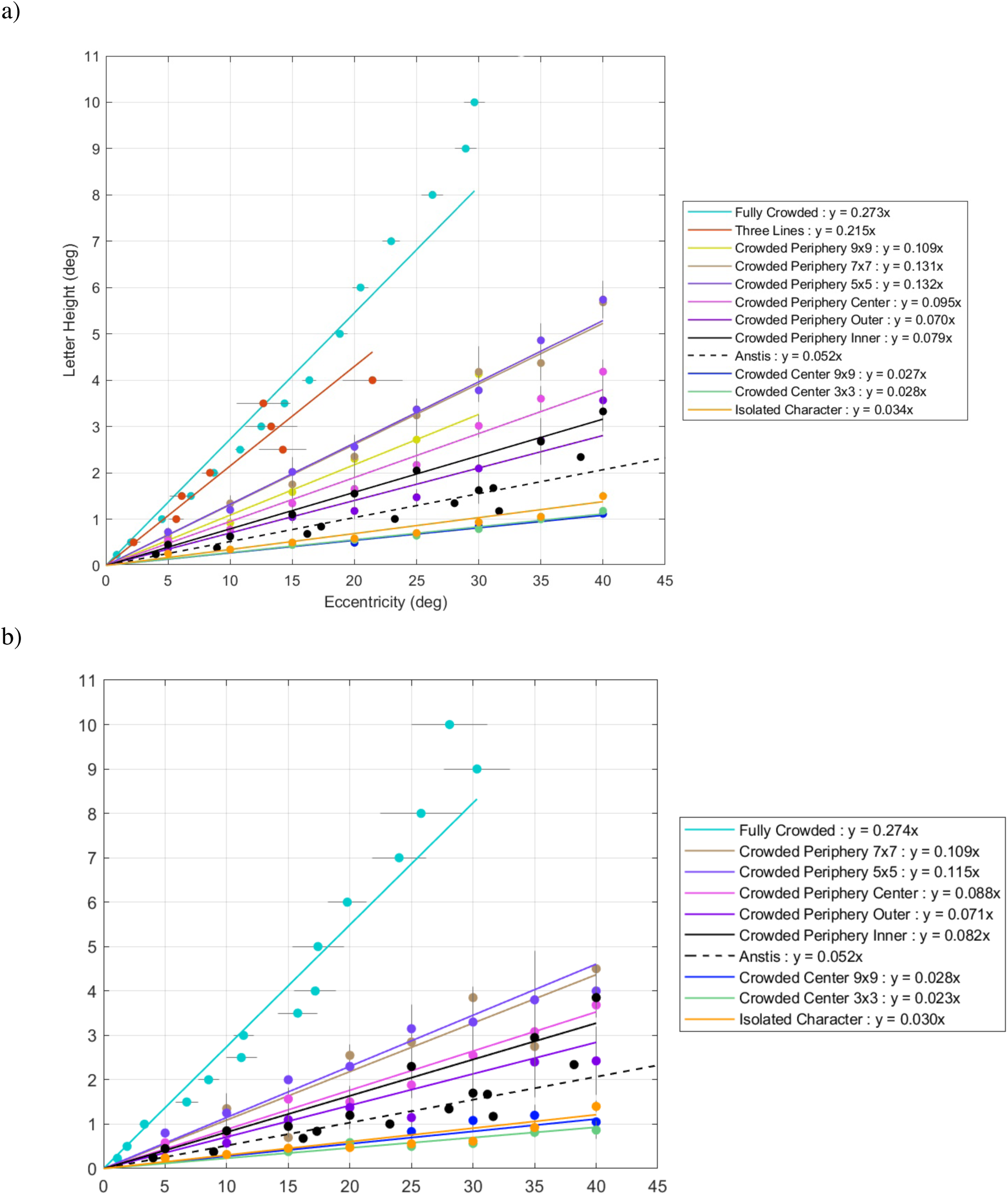
Visual acuity measured by threshold letter height vs. retinal eccentricity for all protocols. Threshold letter height observations at each retinal eccentricity were averaged together to form a single point on the graph. A line of best fit was calculated for each protocol via *χ*^2^ minimization. **a**. Threshold letter height vs. eccentricity results with linear best fit line, for all visual acuity paradigms; results for each eccentricity of each paradigm are averaged across all subjects. **b**. Single subject results of letter height vs. eccentricity across all visual crowding paradigms; this single-subject example confirms that trends were not only observed across subjects but also for individual subjects.

A smaller slope value indicates a better visual acuity. These results show a clear relationship between letter height and eccentricity as a function of crowding (Figure 4). The slopes of both Crowded Center protocols and the Isolated Character protocol were smaller than the slope of data gathered from Anstis’ study (1974). In between the fit for Anstis’ protocol and Fully Crowded protocol, the Crowded Periphery Outer had a smaller slope than those of the Crowded Periphery Center protocols. The visual acuity of the Three Lines protocol falls in between the visual acuity of the Crowded Periphery protocols and the Fully Crowded protocol.

**Figure 4.**
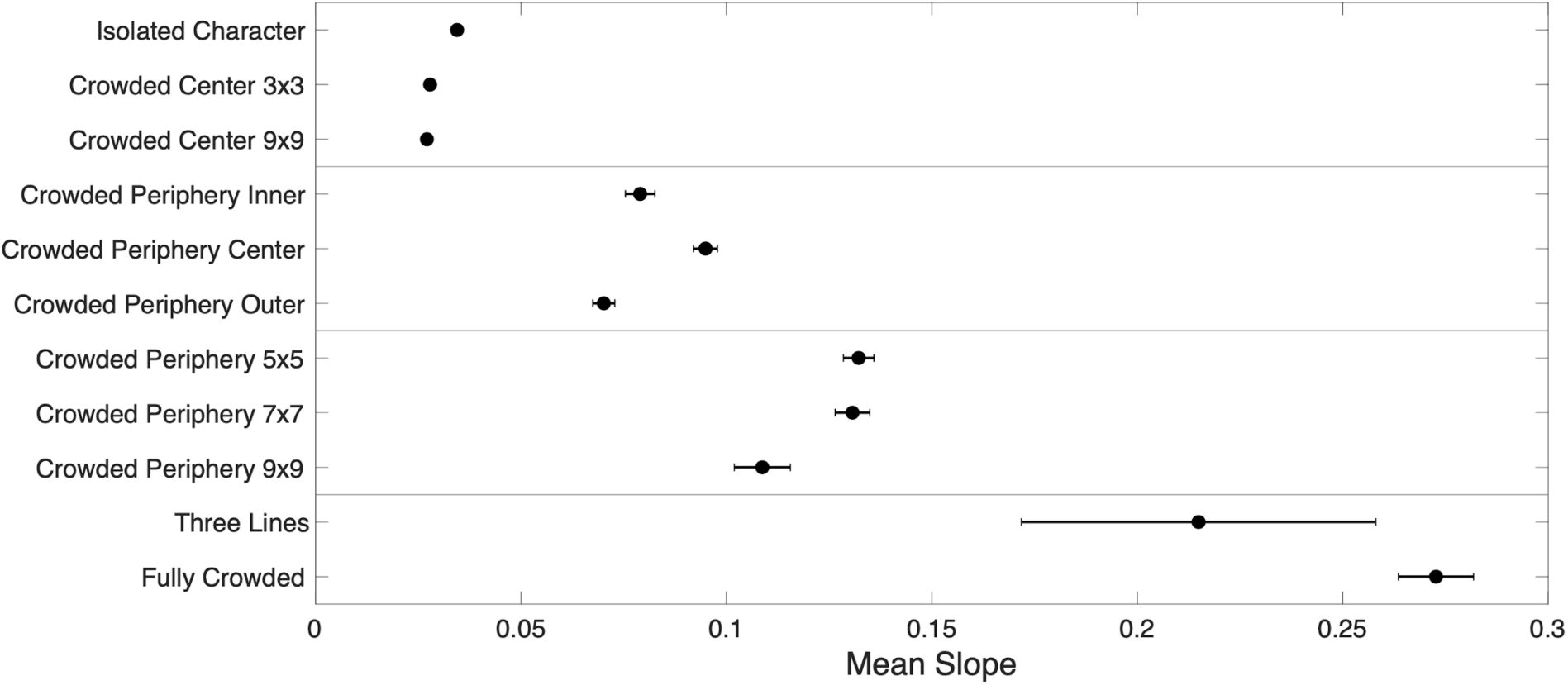
Comparison of normalized visual acuity measured by mean slope of threshold letter height vs. retinal eccentricity for all protocols (i.e. the best fit slope value results shown in Figure 3a). Threshold letter height observations at each retinal eccentricity were averaged together for a single value of threshold letter height for each experimental eccentricity value, across all trials for all participants. Mean slope is defined as the optimized slope, best fit linear free parameter, calculated for each protocol via *χ*^2^ (Chi Square) minimization of the line y = a x. Isolated Character, Crowded Center 3×3, and Crowded Center 9×9 mean slopes: 0.0344 ± 0.0005, 0.0278 ± 0.0007, and 0.0271 ± 0.0009, respectively. Crowded Periphery Inner, Crowded Periphery Center, and Crowded Periphery Outer mean slopes: 0.0790 ± 0.0036, 0.0949 ± 0.0029, and 0.0701 ± 0.007, respectively. Crowded Periphery 5×5, Crowded Periphery 7×7, and Crowded Periphery 9×9 mean slopes are 0.132 ± 0.004, 0.131 ± 0.005, and 0.109 ± 0.007, respectively. Three Lines and Fully Crowded mean slopes were 0.215 ± 0.043 and 0.273 ± 0.109, respectively. Information on how the Standard Error of the slope parameter was extracted can be found in the Statistics Analysis Addendum of the Appendix.

Critically, we determined a statistical significance between Fully Crowded and Crowded Periphery 9×9, Crowded Periphery Outer and Crowded Periphery Center, Isolated Character and Crowded Center 3×3, and Isolated Character and Crowded Center 9×9. The effect of Crowded Periphery Inner vs. Crowded Periphery Outer nearly reached significance. No significant effect was observed between Crowded Center 3×3 and Crowded Center 9×9 (Appendix Table 4).

### Protocol Correlations

The slope for a given behavioral protocol of each recruited subject was plotted against the slope of that subject’s Isolated Character protocol slope (Figure 5). Since all subjects performed the Isolated Character protocol, each protocol was plotted against the subject’s mean slope for the Isolated Character protocol.

**Figure 5.**
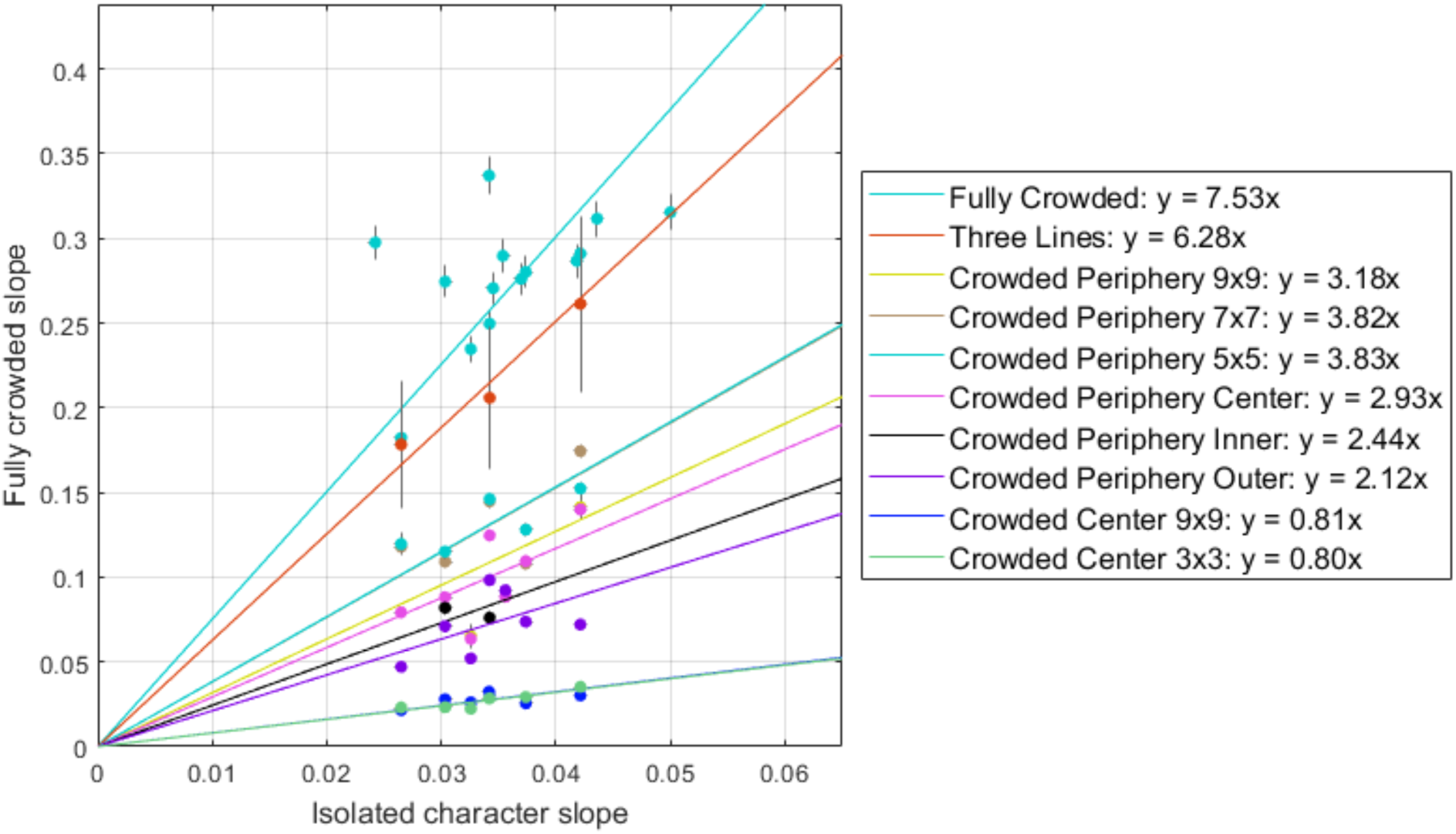
Correlation between slope values of subjects for a given protocol against mean slope of their Isolated Character protocol data. All data was compared against the Isolated Character protocol since every subject performed this protocol. Each data point represents a subject’s slope for a particular paradigm, calculated via *χ*^2^minimization (Figure 3b), plotted against their Isolated Character slope. A line of best fit was calculated via *χ*^2^ minimization. All datasets from a single subject for each protocol were averaged into one data point on a given graph.

These plots showcase the subject-to-subject variation in visual acuity for each protocol; in other words, these plots provide insight into whether subjects who performed well on a given protocol also performed well on the other protocols they completed. A summarized table of these slopes and correlation coefficients is shown below (Table 4).

**Table 3.** The normalized distribution of threshold letter height at given retinal eccentricities (Appendix Fig. 4b) was aggregated across all subjects for each paradigm. A two-tailed Wilcoxon rank sum test was then used to assess the significance between the visual acuities resulting from each paradigm. The p-values resulting from those tests are shown in this table. In each pairwise comparison, the paradigm for which better visual acuity was observed is underlined. The comparisons between Crowded Periphery Inner vs. Crowded Periphery Center and Crowded Periphery Inner vs. Crowded Periphery Outer nearly reached significance. The comparison between Crowded Center 3×3 and Crowded Center 9×9 did not reach significance.

| ***Paradigm vs. Paradigm*** | ***p-value*** |
| --- | --- |
| Fully Crowded vs. <u>Isolated Character</u> | $p < 1E-5$ |
| Fully Crowded vs. <u>Crowded Periphery 9x9</u> | $p < 1E-5$ |
| <u>Crowded Periphery Inner</u> vs. Crowded Periphery Center | $p = 0.011$ |
| <u>Crowded Periphery Outer</u> vs. Crowded Periphery Center | $p < 1E-5$ |
| Crowded Periphery Inner vs. <u>Crowded Periphery Outer</u> | $p = 0.079$ |
| Isolated Character vs. <u>Crowded Center 3x3</u> | $p < 1E-5$ |
| Isolated Character vs. <u>Crowded Center 9x9</u> | $p < 1E-5$ |
| <u>Crowded Center 3x3</u> vs. Crowded Center 9x9 | $p = 0.38$ |

**Table 4.** Summary of parameters shown in Figure 5. Slope parameters and error estimations were calculated via *χ*^2^ minimization. The standard error of the slope parameters was evaluated from the *χ*^2^ contour around the minimum, at values of *χ*^2^_*min*_+ 1.

| *Visual Acuity Paradigm* | **Correlation Parameters** |  |
| --- | --- | --- |
|  | *Slope* | *Correlation Coefficient* |
| Fully Crowded (FC) | $7.53 \pm 0.04$ | 0.54 |
| Three Lines (3L) | $6.28 \pm 0.37$ | 0.98 |
| **Crowded Center (CC)** |  |  |
| Crowded Center 3x3 (CC3) | $0.80 \pm 0.01$ | 0.91 |
| Crowded Center 9x9 (CC9) | $0.81 \pm 0.01$ | 0.57 |
| **Crowded Periphery (CP)** |  |  |
| Crowded Periphery Inner (CPI) | $2.44 \pm 0.05$ | |
| Crowded Periphery Center (CPC) | $2.93 \pm 0.02$ | 0.74 |
| Crowded Periphery Outer (CPO) | $2.12 \pm 0.02$ | 0.46 |
| Crowded Periphery 5x5 (CP5) | $3.83 \pm 0.03$ | 0.78 |
| Crowded Periphery 7x7 (CP7) | $3.82 \pm 0.03$ | 0.66 |
| Crowded Periphery 9x9 (CP9) | $3.18 \pm 0.06$ | 0.40 |

From these results, it is important to note that there exists a spectrum of visual acuity: crowding is not a function simply of the spacing between characters; instead, these results demonstrate that the type and the extent of crowding also impact visual acuity.

The raw data collected from these experiments is publicly available on GitHub (Wilson, 2020).

## Discussion

The results from our behavioral protocols motivate a more comprehensive theory to answer the question: what exactly is the mechanism behind the semantic extraction of peripheral visual information? Previous theory, for example, fails to explain the observed reduction in PVA––one that monotonically decreases with the degree of environmental crowding in the vicinity of our attention. More specifically, previous theory takes a passive approach, arguing that the increase in receptor field size towards the periphery blurs the intake of visual information in that region. However, if this were the only factor impacting PVA, we would not have observed a statistical difference between the mean slopes of the Fully Crowded and Crowded Periphery 9×9 protocols or between the mean slopes of the Isolated Character and the Crowded Center protocols. In both cases, the protocols have the same degree of crowding––and therefore distortion– –in the peripheral region of interest.

These distinct results instead demonstrate the need for a more encompassing theory that is driven by attention and utilizes the dorsal pathway in conjunction with the ventral pathway to extract semantic information from the periphery. We hypothesize that semantic recognition ultimately relies on the notion of coincidence detection: for each object, we have a prior memory or prediction that must coincide with the incoming image that is mapped onto the ventral pathway. This coincidence detection mechanism ultimately makes use of Hebbian plasticity (Hebb, 1949). Semantic recognition is most accurate at the fovea because this image is directly mapped onto V1 after passing through the lateral geniculate nucleus of the thalamus. Implicit in this assumption, however, is that the prior image is mapped in the log-polar coordinate system. The scale invariance of the log-polar system of V1 facilitates rescaling of visual input from the center gaze via translation in order to match the predicted pattern in this coordinate system.

However, as previously noted, this neural pattern present on V1 with the center gaze is significantly distorted towards the periphery. Thus, while the ventral visual pathway can be used to identify objects through scaling invariant perception, information from the dorsal visual pathway is more important for localizing objects (Spering & Carrasco, 2015). Specifically, we hypothesize that the covert attention vector– –driven by the attentional centers of the brain in the dorsal stream––helps identify the center of the peripheral character of interest. Through parallel translation, these dorsal structures are able to internally translate the character to the center where it is mapped in a log-polar manner for semantic extraction through the ventral pathway.

In our results, we observed a continuous reduction in visual acuity spanning from the Crowded Center to the Fully Crowded protocols based on the amount of the crowding surrounding the stimulus character. This supports our model because the neural site that primes the stimulus character to be translated is located in the dorsal pathway. These structures have a resolution that is significantly reduced relative to the resolution of V1 in the ventral pathway; thus, increased crowding distorts the ability of these dorsal structures to identify the center of the stimulus character to be translated internally (Vanni et al., 2015). This alters the image that is ultimately mapped onto the ventral pathway, reducing the accuracy of semantic extraction of the peripheral object in log-polar space.

Prior research supports the concept of an internal shift between systems as a means of coordinating the activity of space distributed over a series of cells (Andersen et al., 1993). While Andersen’s work discusses the coordination between the coordinate system of the visual pathway system and that of the muscular system, we instead focus on the visual pathway specifically. We hypothesize that the parallel translation of the dorsal pathway is activated on the same clock as the ventral pathway for semantic extraction; thus, through the synchronization of these clocks, the dorsal and ventral visual streams can work together to achieve accurate semantic extraction from the periphery.

Previous literature demonstrates that it is reasonable to assume that the coordination of any spatially-distributed information requires traveling waves, and that coordinate shifts in our model could be mediated by a phase shift of these traveling waves. It is also reasonable to assume that there is communication between the different regions responsible for these computations (i.e. translations and transformations) via synchronous oscillations. This is because we already know that brainwaves play a large role in how spatial information is scanned and encoded. For example, the 10 Hz alpha wave operates in a manner analogous to a radar beam scanning different visual hemifields consistent with phase shifts (Lozano-Soldevilla & VanRullen, 2019). Spatial processing also occurs in a non-continuous manner with oscillations of perceptual sensitivity between high periods of attention and diminished periods of attention modulated by the theta phase (Fiebelkorn & Kastner, 2019; Fiebelkorn, Pinsk, & Kastner, 2018). Our model builds upon this previous work––with regards to both crowding impact and oscillations within the brain––proposing a more holistic explanation of how semantic information is extracted in the periphery.

Just as important, we observed that the visual acuity from the Crowded Periphery Inner paradigm was worse than the visual acuity of the Crowded Periphery Outer protocols paradigm (p = 0.079). This directly contradicts previous literature that assumes that, under the same crowding conditions, visual search gets increasingly worse as the character moves from the center to the periphery (Carrasco et al., 1995; Wolfe, O’Neill, & Bennett., 1998). However, the difference observed between visual acuity in the Crowded Periphery Inner and Crowded Periphery Outer results supports our theory which hypothesizes that the attentional vector––much like the saccadic vector motivated by the pulvinar nucleus and the superior colliculus for overt attention––overshoots past the target letter in attempting to localize its center (Bahill, Clark, & Stark, 1975; Kapoula & Robinson, 1986). Thus, despite the same amount of crowding in the receptive field distorting peripheral vision, the visual acuity resulting from the Crowded Periphery Outer paradigm is better than the visual acuity resulting from the Crowded Periphery Inner paradigm. This overshoot of the attention vector centers on the outer character, instead of the middle character, and it is ultimately the scaling invariant pattern of the outer character that is translated and mapped onto the ventral pathway for semantic extraction.

Importantly, a significant difference was observed between stimuli displayed in the left vs. the right visual field in the Isolated Character experiment (p = 8.9E-3). Visual acuity was better for stimuli displayed in the right visual field, consistent with previous research. We hypothesize that this effect may be due to hemispheric differences in the processing of visual stimuli. It has been suggested that the left hemisphere (receiving information from the right visual field) is optimized for identification of the local characteristics of visual stimuli (i.e. identity of a character) (Hugdahl, 2013). Other researchers have suggested that the visual word form area, lateralized to the left hemisphere, is responsible for the improved visual acuity for characters displayed in the right visual field (Hannagan et al., 2015).

Overall, our behavioral protocols demonstrated three novel findings that cannot be explained by current theory: (1) Crowded Center PVA is significantly better than Isolated Character PVA (p < 1E-5), (2) Crowded Periphery 9×9 PVA is significantly better than Fully Crowded PVA (p < 1E-5), and (3) Crowded Periphery Outer PVA is better than Crowded Periphery Inner PVA (p < 0.079). In each of these cases, the amount of crowding in the receptive field of the peripheral target was the same. Thus, we hypothesize a broader, more encompassing theory that utilizes the attention vector to allow the dorsal structures of the visual stream to accurately locate the center of peripheral character. This peripheral character is internally translated and ultimately mapped in the log-polar coordinate system for semantic extraction.

### Future Work

In order to continue assessing our model’s validity, it would be important to utilize eye-tracking to hold subjects more accountable to our behavioral exercises. While we are confident in our subjects’ results because they match the pilot data (i.e. internal and mock-session data) taken by lab members who strictly adhered to the protocol guidelines, it would be helpful to have an external control to ensure subjects are always staring at the center of the display (i.e. at the green dot or arrow). Additionally, the use of eye-tracking would be constructive in determining where exactly the fovea is centered when a subject is examining a character at 0°. This may provide further insight into the mechanisms behind semantic extraction in the periphery, since the neuronal pattern mapped onto the V1 differs depending on the focus of the fovea (Appendix Figure 3).

In the datasets analyzed for this paper, several pilot subjects had taken data for different protocols in different sittings (i.e. a different study session later in the day or on another day entirely). To better control for the variability within each subject’s dataset, it would be important for the subject to collect data in the same sitting for the pairwise comparisons between the mean slope of the Isolated Character protocol and another protocol.

Further, EEG measurements may be able to provide greater insight into the specific brain structures responsible for the semantic extraction of visual information from the periphery. These measurements could track the rhythms (i.e. phase shifts) of cortical waves in various regions of the brain and their correlation to our observed behavioral results.

Overall, our results shed light on the need for a significantly more holistic theory of visual information processing––one that has been hinted at but not yet laid out in its entirety. By examining the entire region of visual space, we offer a plausible, albeit novel, explanation that makes use of both the dorsal and the ventral visual pathways of the brain.

## Acknowledgments

We are grateful to Aaron Blaisdell, Agatha Lenartowicz, Sandra Loo, Zahra Aghajan, Javier Carmona, Chandan Kittur, Chris Dao, Justin Delfino Yi, Tomoyo Namigato, Sonia Niagara Chung, and Trevor McCarthy for their continued support and assistance.

We would like to thank Aliza Ajmal, Charles E. Ang, Mikaila Ann Bantugan, Alex Chang, Edwin Chau, Athena Dong, Wenjie Dong, Emerald Ellspermann, Valeriia Filippska, Zhaochong Han, Marzia Hazara, Sarah Hong, Rachel Houseworth, Rucha Kulkarni, Kenny Le, Sophia R. Luzzi, Cianan Murphy, Natalie Nguyen, Kristina Pagano, Sumaarg Pandya, Preny Riganian, Gianna Tunzi, and Aijun Zhang who contributed to this project during its preliminary stages.

## Appendix

**Appendix Figure 1.**
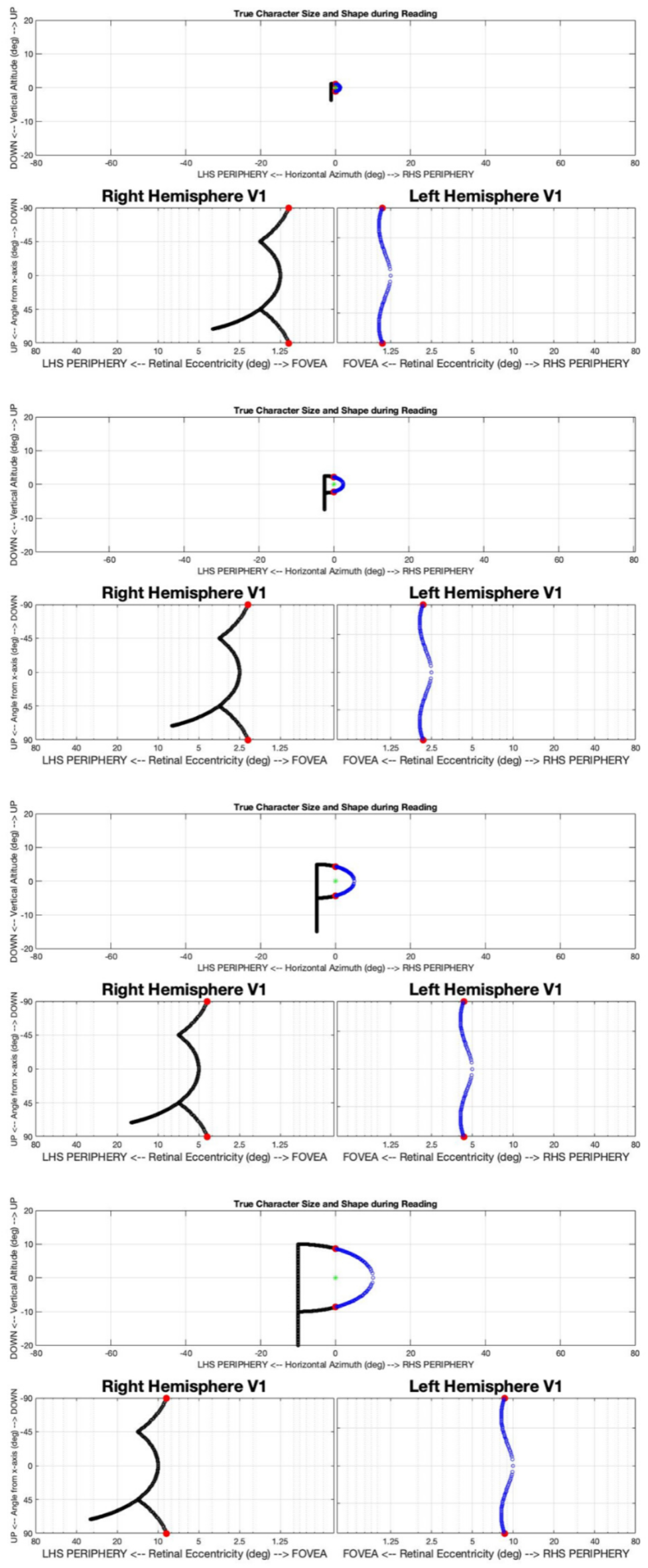
Scaling invariant perception on the visual cortex results in a translation of the character “P” as it is scaled up in visual space.

**Appendix Figure 2.**
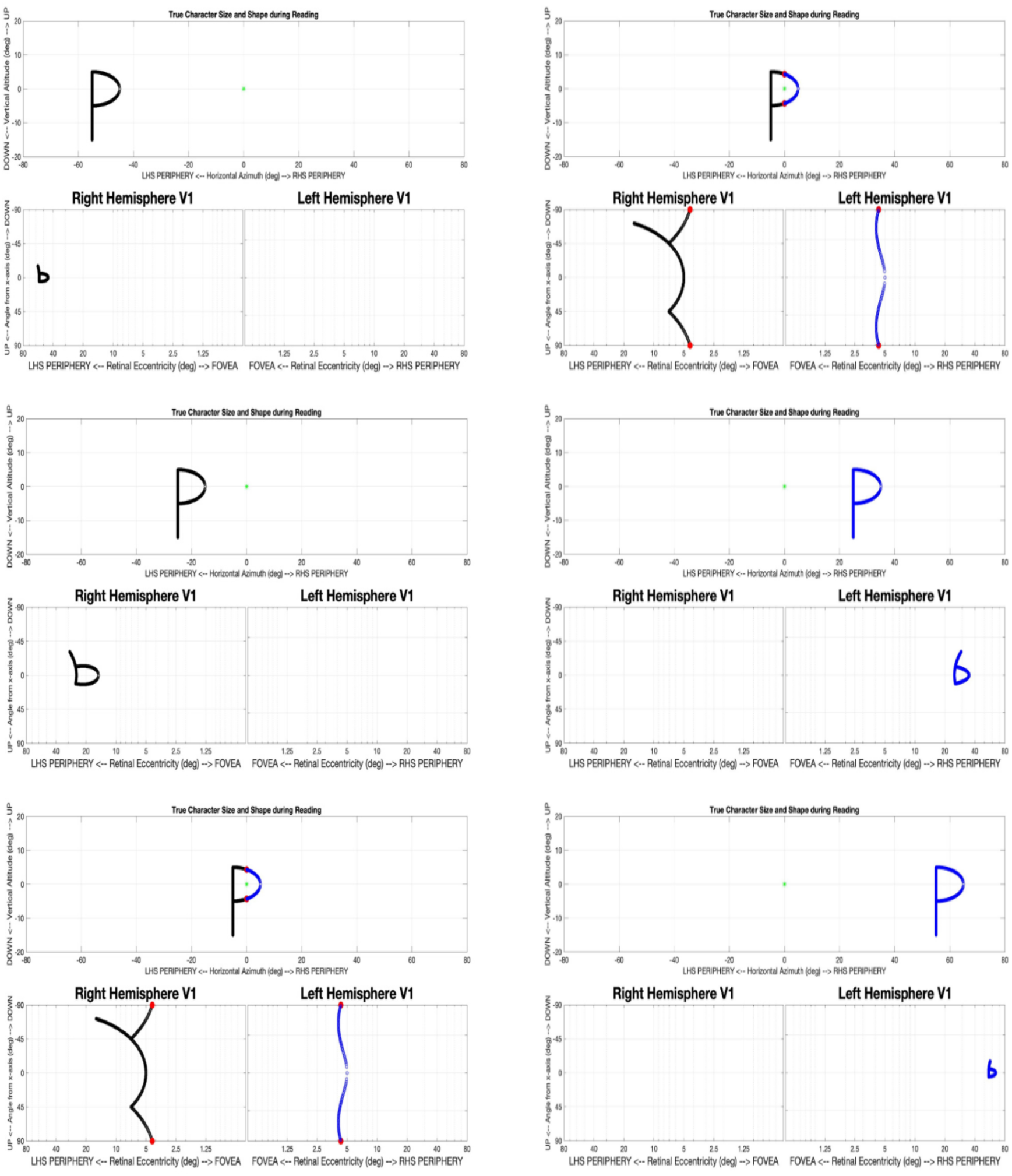
Distortion of the character “P” on the visual cortex when in the periphery. This phenomenon seemingly poses a problem for semantic extraction of peripheral visual information since the brain has been trained to recognize the pattern when it is presented only in center vision.

**Appendix Figure 3.**
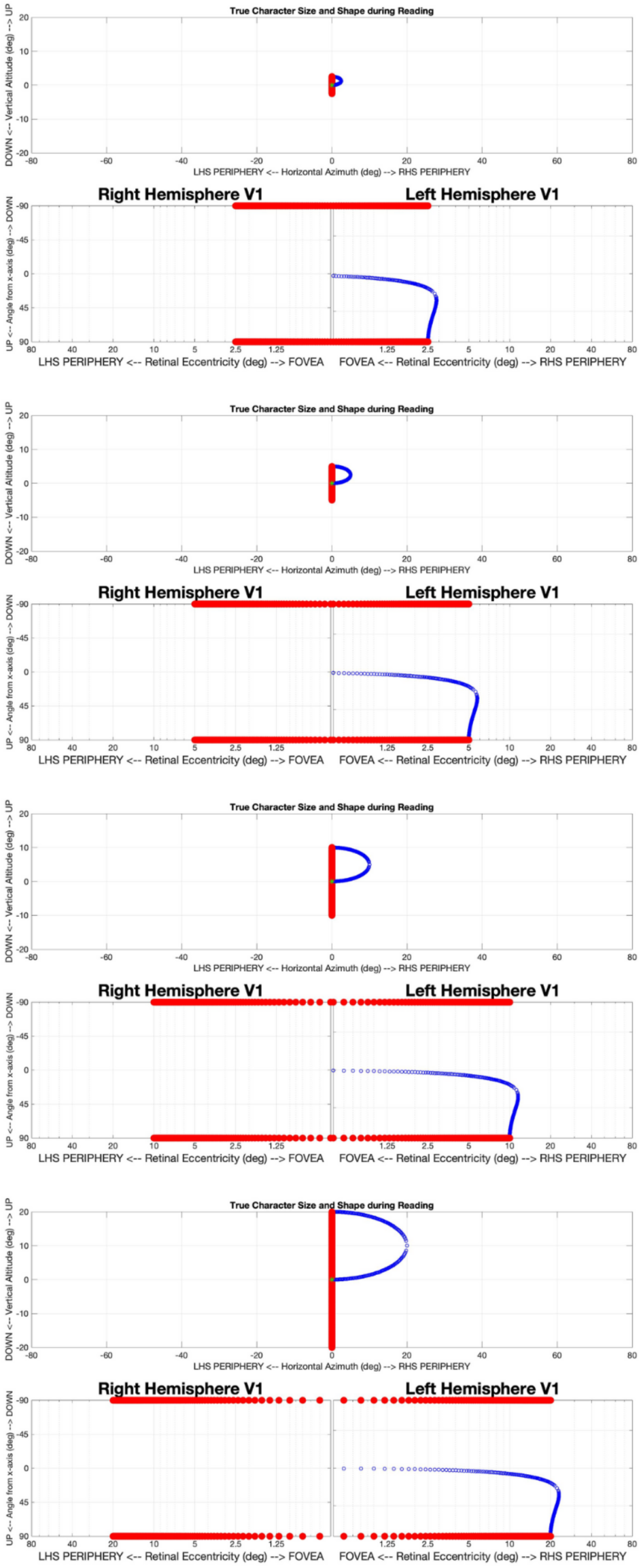
The exact location of the visual gaze, upon an image affects how the figure is mapped onto the visual cortex, independent of the object’s exact location in visual space. In both Appendix Figure 1 and Appendix Figure 3, the focal point of the fovea is denoted as the green dot. In Appendix Figure 1, the fovea is focused on the center of the empty space within the blue “P” loop; in this figure, however, the fovea is focused at the intersection of the vertical red line and the blue “P” loop.

## Statistical Analysis

Threshold letter height observations were normalized by their retinal eccentricity values,

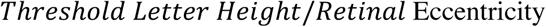

which produced a roughly normal distribution (Appendix Fig. 4b). Outliers, defined as points which deviated from the mean of this distribution by more than 2.5 *σ*, were recursively removed from the data set and assumed to be the result of subject error. For example, in the case of the Fully Crowded protocol, outliers most often took the form of small eccentricity observations recorded at large letter heights. An observation such as this, which deviated so significantly from the rest of the distribution, could have been the result of the subject accidentally pressing the incorrect button when identifying the character. Observations made at retinal eccentricities in the range of 5° to 15° were excluded from the Crowded Center protocols, as it is impossible to guarantee that crowding effects due to the target character’s proximity to the central block of text did not worsen visual acuity.

**Appendix Figure 4.**
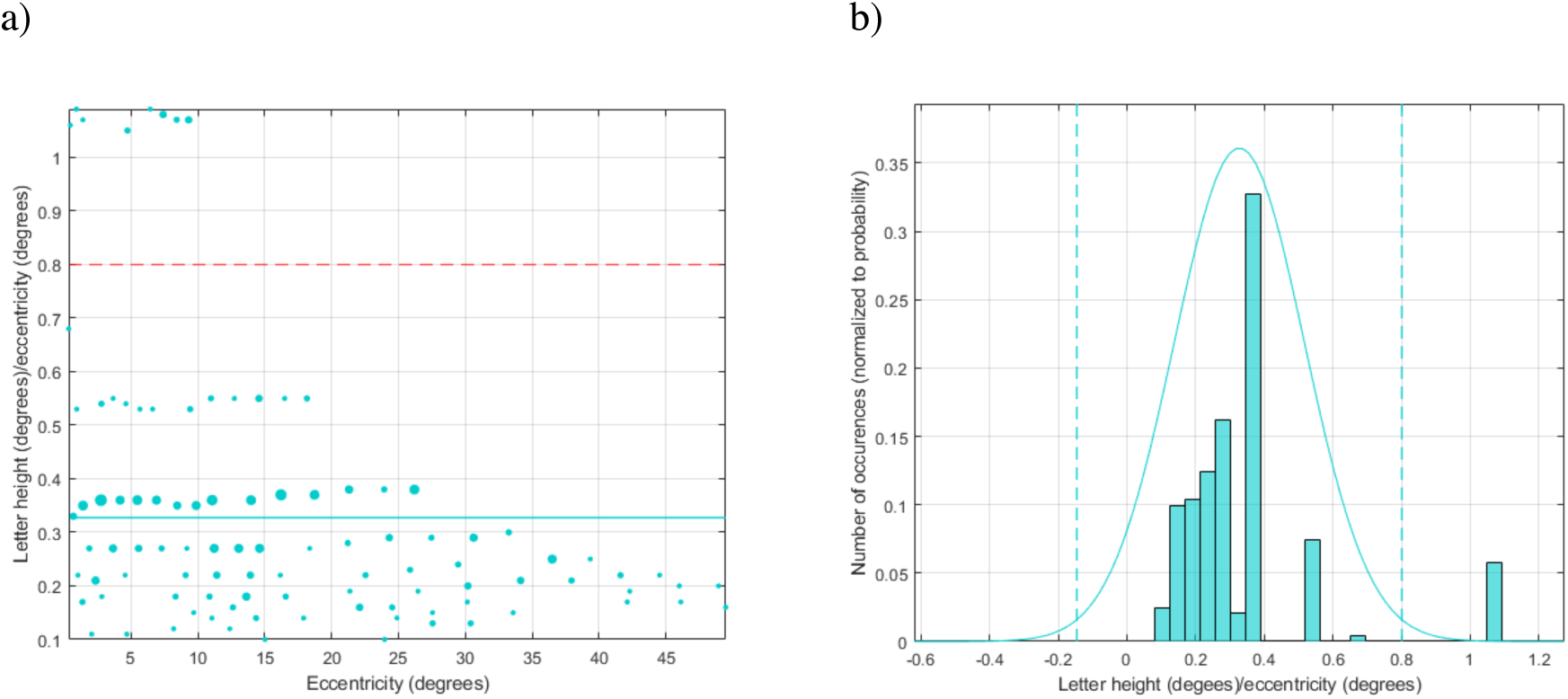
Graphs of a single subject’s Fully Crowded normalized threshold letter height. (a) Normalized threshold letter height vs. retinal eccentricity. A horizontal line of best fit of the form *y* = *a* represents the mean of the distribution. A dashed red line demarcates a value of +2.5*σ* from the mean. (b) Histogram with a gaussian fit of the distribution of normalized threshold letter height values. Dashed vertical lines represent values of ±2.5*σ* from the mean.

This truncated set of normalized data was then transformed back into the linear scale and replicate observations at each discrete measurement value were averaged. A line of best fit fixed at the origin (Figure 3), and a line of best fit with a free intercept parameter were calculated by minimizing the *χ*^2^,

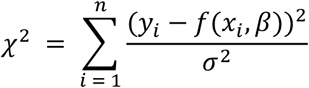

where *y*_*i*_ represents the observed value, *σ*^2^ represents the square of the estimated standard error of the observation, and the equation *f*(*x*_*i*_, *β*) is of the form *y*_*e*_ = *β* _1_*x* + *β* _2_. The standard error of each observation was estimated as the standard error of the distribution of every measurement taken at that discrete value, across all subjects.

For both fits, *χ*^2^ vs. *β* was plotted (Appendix Figure 5). The standard error of the fit parameters was evaluated from the *χ*^2^ contour around the minimum, at values of *χ*^2^_*min*_ + 1.

**Appendix Figure 5.**
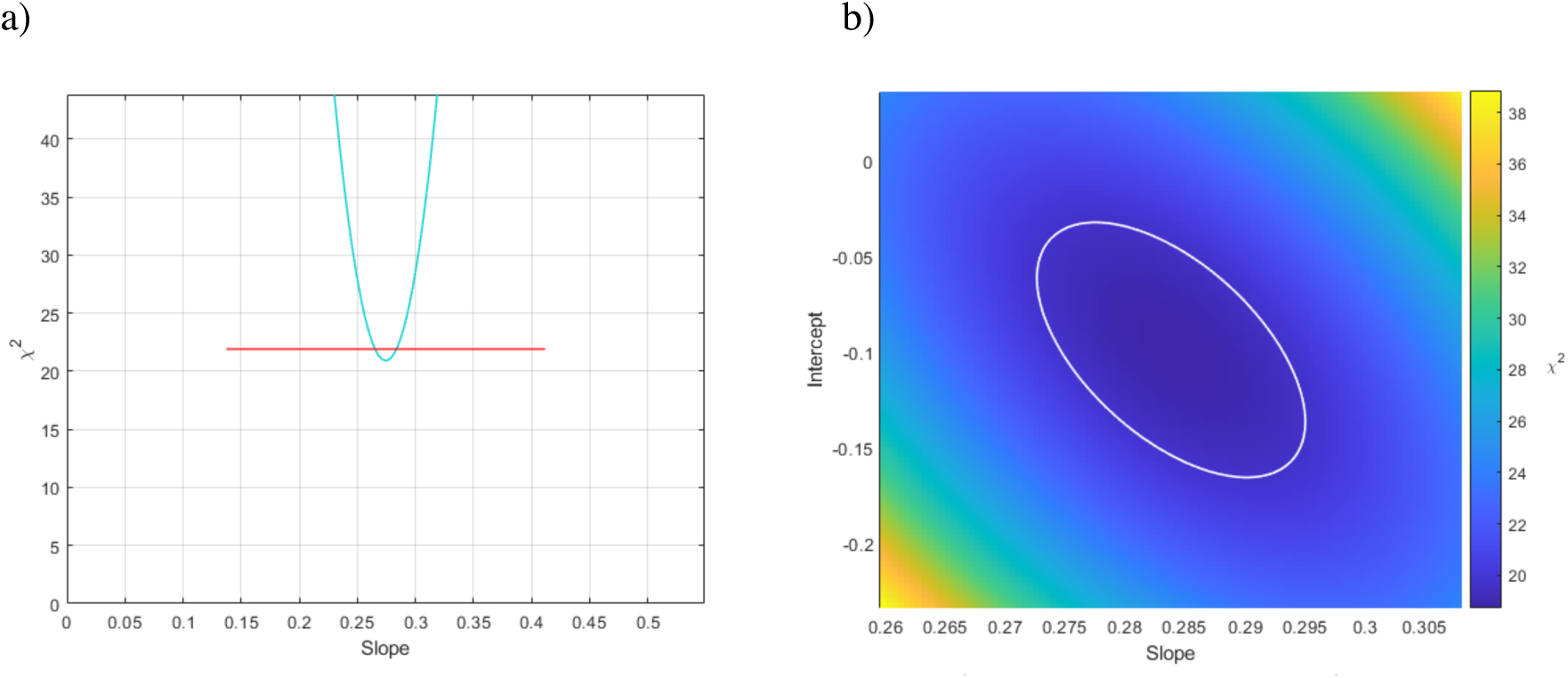
Graphs of a single subject’s Fully Crowded *χ*^2^ vs. fit parameters. (a) *χ*^2^ vs. slope parameter for the *y* = *ax* fit, displaying a parabolic curve at the minimum. A red horizontal line demarcates a *χ*^2^ value of *χ*^2^ _min_+ 1. (b) *χ*^2^ vs. slope and intercept parameter for the y = *a*x + *b* fit. A white oval demarcates a *χ*^2^ value of *χ*^2^_min_ + 1.

The fit without a free intercept parameter was desirable in order to easily evaluate correlation between a subject’s performance on different protocols. In order to validate this fit, data from all subjects was combined and optimal fit parameters were calculated with and without a free intercept parameter. The *χ*^2^_*ν*_(*χ*^2^_*min*_*/degrees of freedom*) of the fixed intercept fit was sufficiently close to 1 across all protocols, and no significant improvement was seen with a free intercept parameter (Appendix Table 1). Results were plotted against those of Anstis (1974). The data points from Anstis’ paper were extracted by using plot digitizer software (Huwaldt & Steinhorst, 2015).

**Appendix Table 1.** *χ*^2^_*ν*_ for y = ax and y = ax + b fit of data averaged across all subjects. *χ*^2^_*ν*_ is the *χ*^2^_*min*_ value divided by the degrees of freedom. The values were deemed sufficiently close to 1, and so a linear best fit of only one free parameter, y = *a*x, without y-intercept free parameter, was used for all reported results.

| *Visual Acuity Paradigm* | $y = ax \chi_v^2$ | $y = ax + b \chi_v^2$ |
| --- | --- | --- |
| Fully Crowded | 1.2 | 1.2 |
| Three Lines | 0.081 | 0.094 |
| Crowded Periphery 9x9 | 0.89 | 1.06 |
| Crowded Periphery 7x7 | 0.79 | 0.72 |
| Crowded Periphery 5x5 | 0.57 | 0.67 |
| Crowded Periphery Inner | 1.7 | 1.1 |
| Crowded Periphery Center | 1.8 | 2.04 |
| Crowded Periphery Outer | 1.3 | 1.5 |
| Crowded Center 9x9 | 1.2 | 0.38 |
| Crowded Center 3x3 | 0.74 | 0.86 |
| Isolated Character | 23 | 4.7 |

In order to assess the correlation between a subject’s performance across different protocols, the slope parameter for each protocol was plotted vs. the Isolated Character slope parameter (Figure 4). A line of best fit was calculated for each protocol by *χ*^2^ minimization. The *σ* of each point was estimated as the mean of the subject’s standard deviation of their slope parameter for both of the protocols being assessed. The error bars in the correlation plots represent values of the slope parameter in the negative and positive directions which resulted in a *χ*^2^_*ν*_+1 from the minimum (Appendix Fig. 5a).

Prior to the transformation back into the linear scale, the normalized distribution of observations (Appendix Fig. 4b) was aggregated across all subjects. A two-tailed Wilcoxon rank sum test was used to assess the significance of different protocols. The aggregated distribution from a single protocol was split between direction (i.e. left vs. right visual field measurements) and a two-tailed Wilcoxon rank sum test was again used to assess significance.

MATLAB was used for all statistical analysis. The analysis source code is publicly available on GitHub (Wilson, 2020).

